# Using the UniFrac metric on Whole Genome Shotgun data

**DOI:** 10.1101/2022.01.17.476629

**Authors:** Wei Wei, David Koslicki

## Abstract

The UniFrac metric has proven useful in revealing diversity across metagenomic communities. Due to the phylogeny-based nature of this measurement, UniFrac has historically only been applied to 16S rRNA data. Simultaneously, Whole Genome Shotgun (WGS) metagenomics has been increasingly widely employed and proven to provide more information than 16S data, but a UniFrac-like diversity metric suitable for WGS data has not previously been developed. The main obstacle for UniFrac to be applied directly to WGS data is the absence of phylogenetic distances in the taxonomic relationship derived from WGS data. In this study, we demonstrate a method to overcome this intrinsic difference and compute the UniFrac metric on WGS data by assigning branch lengths to the taxonomic tree obtained from input taxonomic profiles. We conduct a series of experiments to demonstrate that this WGSUniFrac method is comparably robust to traditional 16S UniFrac and is not highly sensitive to branch lengths assignments, be they data-derived or model-prescribed. Code implementing a prototype of WGSUniFrac along with paper reproducible are available at https://github.com/KoslickiLab/WGSUniFrac.

## 1 Introduction

The study of microbial composition and diversity has demonstrated its value in both clinical [5,8,11] and environmental [36] studies. Within-sample diversity (known also as the alpha-diversity) metrics, such as the Shannon index and Simpson diversity, have been used to evaluate and quantify microbial diversity in various settings [19]. In contrast, between-sample (or, the beta-diversity) measurements allow measurement and analysis of differences across multiple samples, giving insights to their significance [16, 48, 49]. Among the most frequently utilized beta-diversity metrics is UniFrac [14, 24, 25, 27, 32, 46].

UniFrac measures the phylogenetic differences between two microbial communities by calculating the fraction of branch lengths unique to one of the two communities on a phylogenetic tree that has been annotated with the predicted abundances of organisms in the two communities [28]. This computation is established on the intuition that the degree to which two communities or environments differ is positively correlated to the degree of difference in the evolutionary path undergone that resulted in the observed divergence: the longer the evolutionary path, the more divergent [26]. Since its introduction in 2005, the UniFrac distance has been widely applied [11,14,48] and its robustness has stood the test of time [27]. Over time, the UniFrac metric has undergone a series of developments ranging from conceptual understanding and application to computation efficiency. The variation of weighted UniFrac was introduced two years after the introduction of the original unweighted version [28]. Fast UniFrac made its debut in 2010, improving the speed of UniFrac computation, hence expanding its application to larger datasets [15]. In 2012, the understanding of the UniFrac distance being equivalent to the earth mover’s distance was brought to light [12], based on which an exact linear-time computation algorithm, EMDUniFrac, was later developed [30] and then later implemented in Striped UniFrac [30]. All these demonstrate the popularity and potential of the UniFrac metric.

In this paper, we discuss the possibility of applying the UniFrac metric to a new type of data: whole genome shotgun metagenomic samples. Traditionally, UniFrac has been employed almost exclusively in the analysis of 16S rRNA sequencing data. The 16S rRNA sequencing method involves amplification and sequencing of the 16S small subunit ribosomal RNA which contains both highly conserved and variable regions, which leads to a simple and cost effective “fingerprinting” approach to inferring microbial composition [41, 42]. An alternative approach to 16S rRNA sequencing is whole genome shotgun sequencing (WGS). Despite requiring more effort and cost, the advantages of WGS analysis are also apparent: higher accuracy, sensitivity, and access to the entirety of the genetic material in a given sample [42]. Additionally, WGS data are becoming more frequently utilized by clinicians and biologists [2, 4] due in part to the ever-decreasing price. Indeed, recent studies have been calling for replacement of 16S rRNA data by WGS data in certain fields of application [4] due to the advantages of WGS.

Though UniFrac is widely employed in the analysis of 16S rRNA and other amplicon studies, it has yet to find its application in WGS metagenomic data. While 16S rRNA and other amplicon sequencing approaches naturally have a single gene to build a phylogeny with, there is no consensus in the metagenomic community on how to best construct a phylogenetic tree from WGS data, with approaches ranging from a variety of single gene approaches [23, 37, 45], whole genome alignment approaches [13, 47], to k-mer based similarity techniques [1, 22, 40]. As such, researchers have primarily focused on utilizing taxonomic trees instead of phylogenetic trees due to the relative ease of identifying taxa present in a sample [34, 38, 44]. Since UniFrac was originally intended for usage on a phylogenetic tree, this difference in underlying tree structure in amplicon studies versus WGS studies explains why UniFrac has not been used in WGS metagenomic analyses. In particular, the absence of phylogenetic relationship among taxa in a taxonomic tree, as well as evolutionary distances reflected in branch lengths, hinders the direct computation of UniFrac. Even so, the robustness of UniFrac demonstrated in numerous amplicon studies motivates the endeavor to overcome this intrinsic difficulty and extend its application to WGS data.

In this paper we demonstrate that by assigning branch lengths to the corresponding taxonomic tree, UniFrac can be applied to WGS data and achieve reasonable robustness. We call this extension WGSUniFrac. A summary of how WGSUniFrac works is shown in Figure 1. Code implementing a prototype of WGSUniFrac along with paper reproducible are available at https://github.com/KoslickiLab/WGSUniFrac.

**Figure 1:**
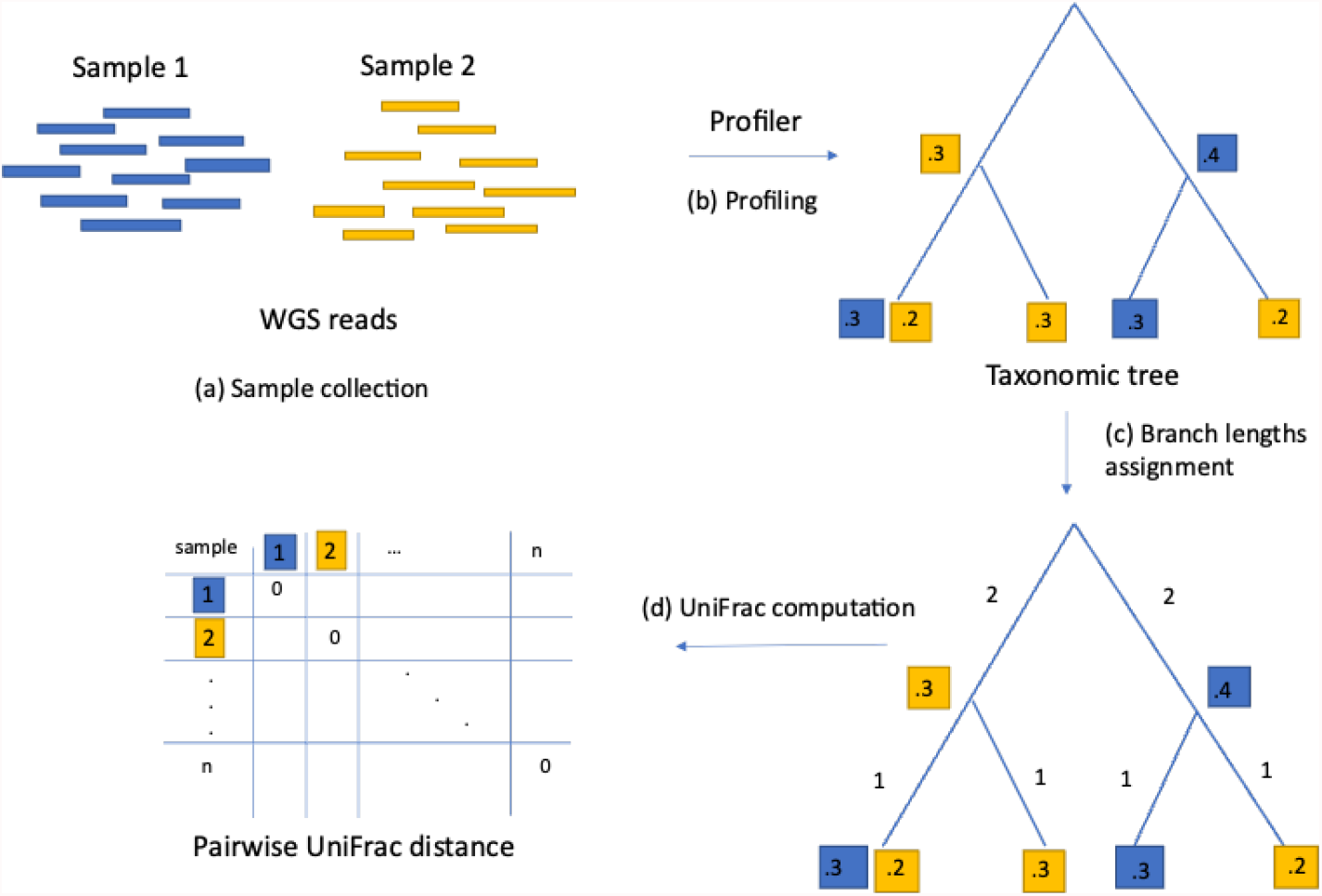
An illustration of the WGSUniFrac workflow. (a) WGS Metagenomic samples are collected. (b) Each sample is converted to its corresponding taxonomic profile using a profiler of choice. Each profile contains the relative abundances of all the organisms present in the sample at all taxonomic levels. The collection of all profiles form a taxonomic tree. (c) Branch lengths are assigned to the taxonomic tree according to branch lengths function specified. (d) Pairwise UniFrac values of all samples are computed using the EMDUniFrac algorithm.

## 2 Results

### 2.1 On taxonomic data converted from phylogenetic data

To test the hypothesis that assigning branch lengths to a taxonomic tree allows computation of UniFrac that reflects beta diversity using only WGS data, we begin with the most ideal scenario: one in which the taxonomic profile of the WGS data exactly reflects the phylogenetic profile of the 16S rRNA data. To this end we constructed the most ideal taxonomic profiles as follows: using the mapping file provided in Greengenes database [10] that maps 16S OTUs with a known phylogenetic tree to their corresponding NCBI taxonomic IDs (taxIDs), we converted a phylogenetic sample to its taxonomic counterpart by simply changing the ID type while maintaining the relative abundance of each species. Using the lineage information associated with the taxID of each species in NCBI, we constructed the full taxonomic profile with the ranks of superkingdom, phylum, class, order, family, genus, and species, representing the taxonomic relations among the species.

Since UniFrac is frequently used to observe qualitative difference in samples when partitioned by certain metadata variables and viewed on a Principal Coordinates Analysis (PCoA) plot, we evaluated the performance of UniFrac computed on such a taxonomic profile based on the hypothesis that if the method makes biological sense, the clustering of samples in the WGS data should agree with that using 16S data. As such, we assessed the performance of WGSUniFrac by observing the clustering of samples under PCoA in comparison to that of their 16S counterparts, as well as quantitatively evaluated the clustering quality with commonly used clustering evaluation metrics.

To better observe the clusters, we created a simple model to mimic samples collected from two distinct environments with the aid of the given phylogenetic tree. To create samples from an environment, we first select a random leaf node on the phylogenetic tree and call it a pivot node. We then randomly selected a fixed number of nodes sufficiently close to the pivot node first selected. To create samples from the other environment, we select a second pivot node sufficiently far away from the first node chosen, and create samples in the same manner centering on the second pivot node. For simplicity of computation, when the distance between two leaf nodes was considered, instead of considering the actual distance in the sense of total branch lengths separating the two nodes, we considered the position of the second node in a list of all nodes ranked according to distance with respect to the first node. For instance, instead of considering “nodes within *x* units of branch length from node 1”, we would consider “nodes among the *y* (for example, 500) nodes closest to node 1”. Throughout this paper, we will call this aforementioned value *y* the “range” of an environment. The distance between the two pivot nodes is also defined in this manner, which we will call “dissimilarity” in this paper (refer to Figure S1). This proxy of replacing the actual distance by the relative position of a node in a list of ranked nodes may very likely result in nonlinearity in the relationship between clustering score and the range or dissimilarity setting, as well as greater variability among repeated experiments having identical range or dissimilarity setting. Nonetheless, it greatly simplifies the calculation and it should not affect the general trend that the greater the dissimilarity and the smaller the range, the more tightly clustered the samples would be on the given phylogenetic tree.

To respectively test the effect of range and dissimilarity on the quality of clustering, we first fixed the dissimilarity to be the maximum (35461) and generated data across ranges 200, 500, 1,000, 5,000, 10,000, 15,000 and 20,000, and then generated data with dissimilarities 800, 900, 1,000, 5,000, 10,000, 20,000, 30,000 and maximum respectively for a fixed range of 500. We generated 100 replicates for each of these setups, each consisting of 25 samples for each environment, with 200 organisms approximately exponentially distributed in relative abundances in each sample. The quality of clustering for each replicate was assessed with the Silhouette Index [43].

In this experiment, the branches of the taxonomic tree were set to the reciprocal of the depth of the branch in the tree (i.e. 1/ distance from root node); we investigate other branch length specifications subsequently. Figure 2 shows the overall results of this experiment, with the trends demonstrated by the plots being expected and intuitive. Namely, the higher the dissimilarity, the greater the differences between samples from the two environments, resulting in more distinguishable clusterings (reflected in higher Silhouette scores). On the other hand, increasing range indirectly decreases dissimilarity by spreading out the clusters/environments, resulting in a decreasing trend of clustering quality. It is noteworthy that these trends were observed in both WGSUniFrac and 16S UniFrac with similar sensitivity. The same trend was observed when other clustering metrics are used (Figure S4). It is also interesting to note that it appears WGSUniFrac is less sensitive to changes in range compared to 16S UniFrac.

**Figure 2:**
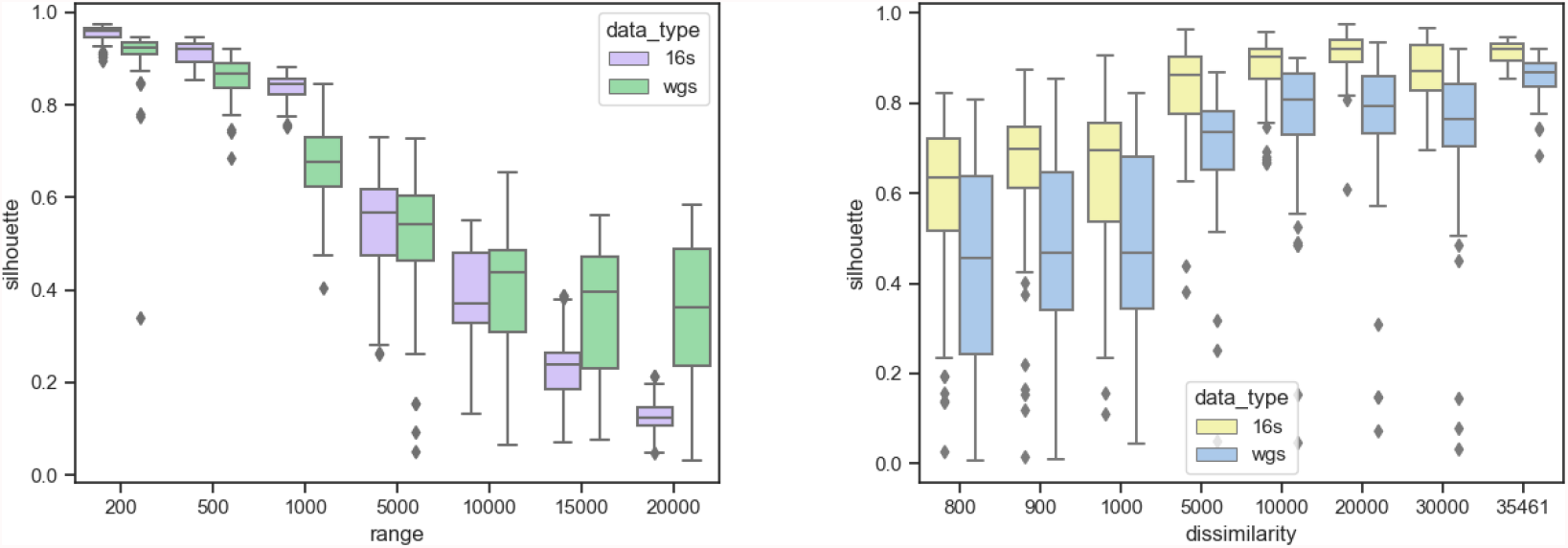
A comparison between the Silhouette scores computed using 16S data and WGS data under different settings of range and dissimilarity. Higher Silhouette score indicates better clustering.

### 2.2 Insensitivity to model or data derived branch length assignment

#### 2.2.1 Model-based branch length assignment

Since the consensus on how branch lengths should be assigned, if it ever exists, has yet to be established, in this section we examine the impact of different branch lengths assignments on WGSUniFrac performance. We first investigated three major categories of branch lengths assignment with respect to the depth of the tree: increasing, constant, decreasing. To this end we defined a branch lengths function to compute the length of a branch located *x* nodes away from the root, denoted by *l*(*x*), by *l*(*x*) = *x*^*k*^ for some integer *k*. In other words, the only factor we take into consideration was the depth of the branch in the tree. We first compared the results by repeating the experiment in the previous section with *k* set to −1, 0, and 1, resulting in decreasing, constant, and increasing branch lengths respectively, when viewed from the root to the leaves.

From Figure 3, the branch length function *x*^−1^ yields the best performance, followed by constant branch length assignment, while assigning branch lengths proportional to tree levels yields the worst result. This is consistent with the observation that organismal similarity increases as one moves to lower taxonomic rank.

**Figure 3:**
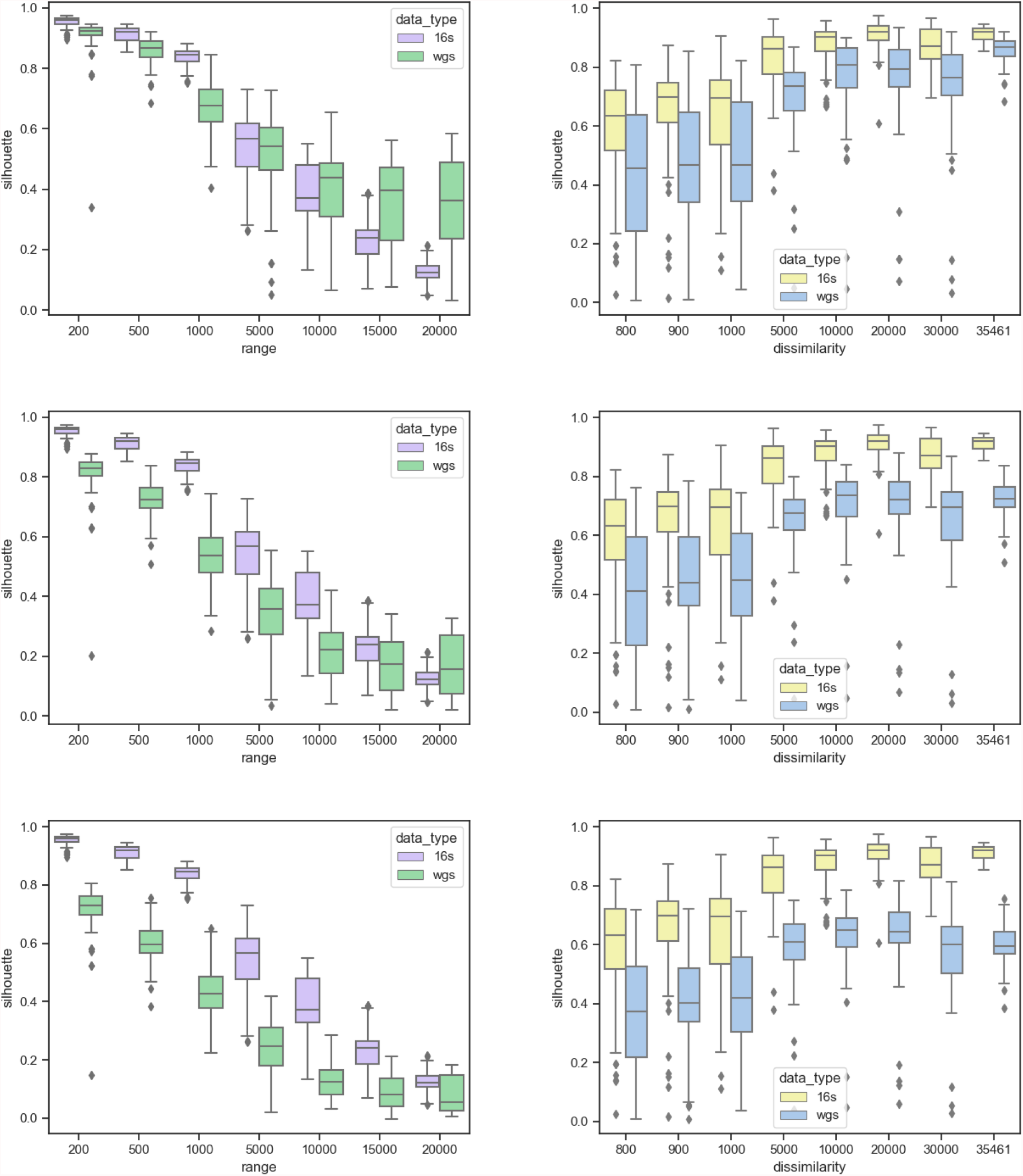
The effect of branch lengths choice. From top to bottom: *k* = −1 (decreasing branch lengths down the tree, *k* = 0 (uniform branch length), *k* = 1 (increasing branch lengths down the tree).

Upon establishing the general relationship between the branch lengths and the depths of the tree, we then examined how sensitive the performance is with respect to fine-tuning of *k* by setting *k* to be -2, -1.5 and -0.5 and repeat the procedure. The results are shown in Supplementary Figure S2, in which we observed an improvement of WGSUniFrac in comparison to the 16S UniFrac with respect to increasing magnitude of *k* (i.e. more negative). This improvement is much more drastic with respect to range than with respect to dissimilarity. In other words, the within-sample diversity is more sensitive to the fine-tuning of ratios between branch lengths. In terms of dissimilarity, which is an intuitive reflection of beta diversity, the improvement in comparison to 16S UniFrac is far less apparent, especially when dissimilarity is small. As such, we conjecture that the magnitude of *k* does not have a significant effect on detecting beta diversity, although it can be suggestive that WGSUniFrac may potentially be more robust than 16S UniFrac when within-sample diversity is large.

For the subsequent experiments, we only considered the branch length function *x*^−1^ in all calculations unless otherwise stated and we revisit the effect of branch lengths selection in Section 2.4 below.

#### 2.2.2 Branch lengths specified with data derived phylogeny-aware taxonomy

In this section, we further examine the robustness of WGSUniFrac with the aid of data obtained from Genome Taxonomy Database (GTDB), a database providing taxonomic trees with topology and branch lengths derived based on protein phylogeny [39]. As a basis of comparison, we used the bac120 tree from GTDB, which is a tree with branch lengths reflecting the phylogenetic information as inferred from the concatenation of 120 marker genes [39].

To assess the impact of branch length specification, we first investigated the performance of WGSUniFrac when the actual branch lengths on the bac120 tree were replaced by the assignment according to the *x*^−1^ function, following the same experimental setup in section 2.1. The results are shown in Figure 4.

**Figure 4:**
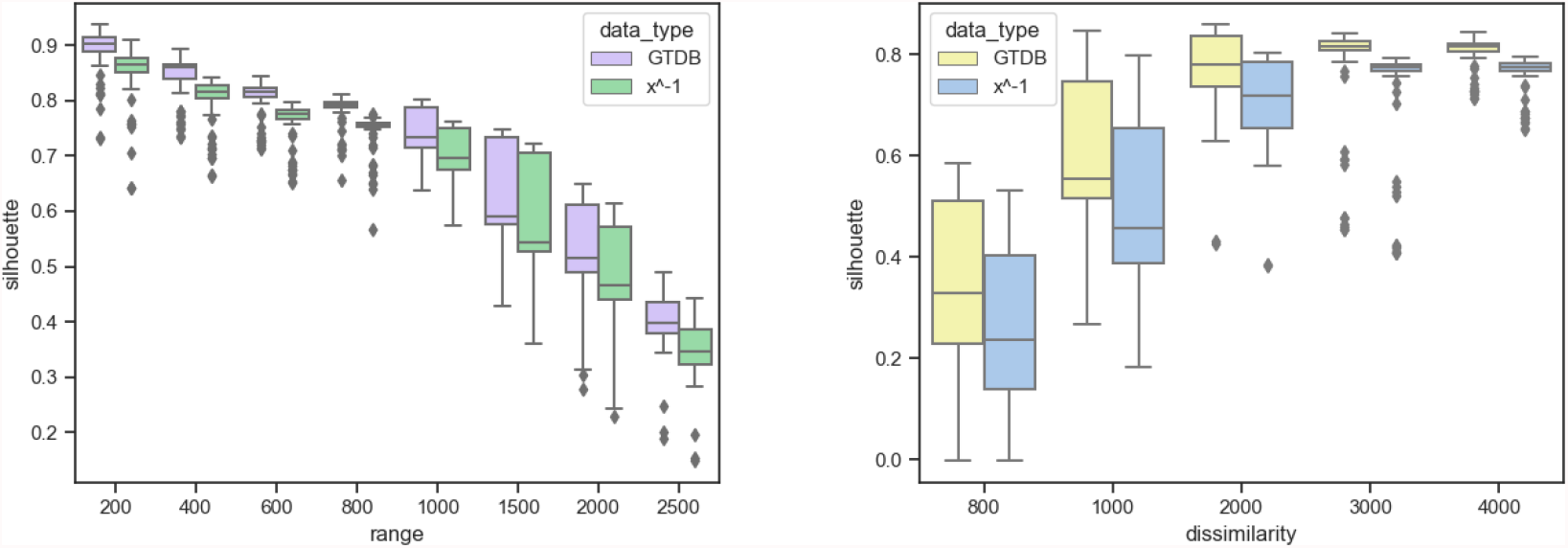
A comparison between the Silhouette scores computed using the GTDB tree and that using the transformed tree with branch lengths reassigned according to branch length function *x*^−1^.

From Figure 4, it can be noted that the behavior of UniFrac computed using the transformed tree closely mimics the original bac120 tree from GTDB, though slightly inferior in all cases. Though a different type of tree was used and different types of data were compared, the nature of this experiment was, in actuality, very similar to that in section 2.1. In both cases, we tested how robust UniFrac would remain when a phylogenetic tree of finely annotated branch lengths was replaced by one that only reflected a general trend instead of having finely labeled branches. The stories told in the two cases were also similar: phylogenetic information does add quality to UniFrac, though UniFrac still reflects general trends without it. In fact, a tree reflecting a general trend among the organisms is sufficient for UniFrac to offer descent insights into beta diversity.

We next investigated the effect of difference in taxonomic topology on UniFrac. According to the authors of GTDB, more than half of the genomes in GTDB had changes in their existing taxonomy [39], resulting in significant differences in the GTDB taxonomy and the existing NCBI taxonomy. As such, among around 4979 organisms having both complete GTDB and NCBI taxonomy, we selected 200 for each sample according to the protocol in section 2.1. For each sample, we generated taxonomic profiles according to GTDB taxonomy and NCBI taxonomy respectively, each having identical organisms and relative abundance distribution. For both taxonomies, we used the branch lengths function *x*^−1^. Fixing dissimilarity to be 4000 nodes apart on the GTDB tree, we created samples with varying values of range, ranging from 200 where nodes from two environments were most tightly clustered, to 2500 where the two environments were slightly overlapping. Similarly, to test the performance under different values of dissimilarity, we fixed range to be 600 and generated samples having dissimilarities ranging from 800, where the two environments were relatively similar, to 4000, where the two environments were highly distinct. Each of these setups was repeated 100 times. The results are shown in Figure 5.

**Figure 5:**
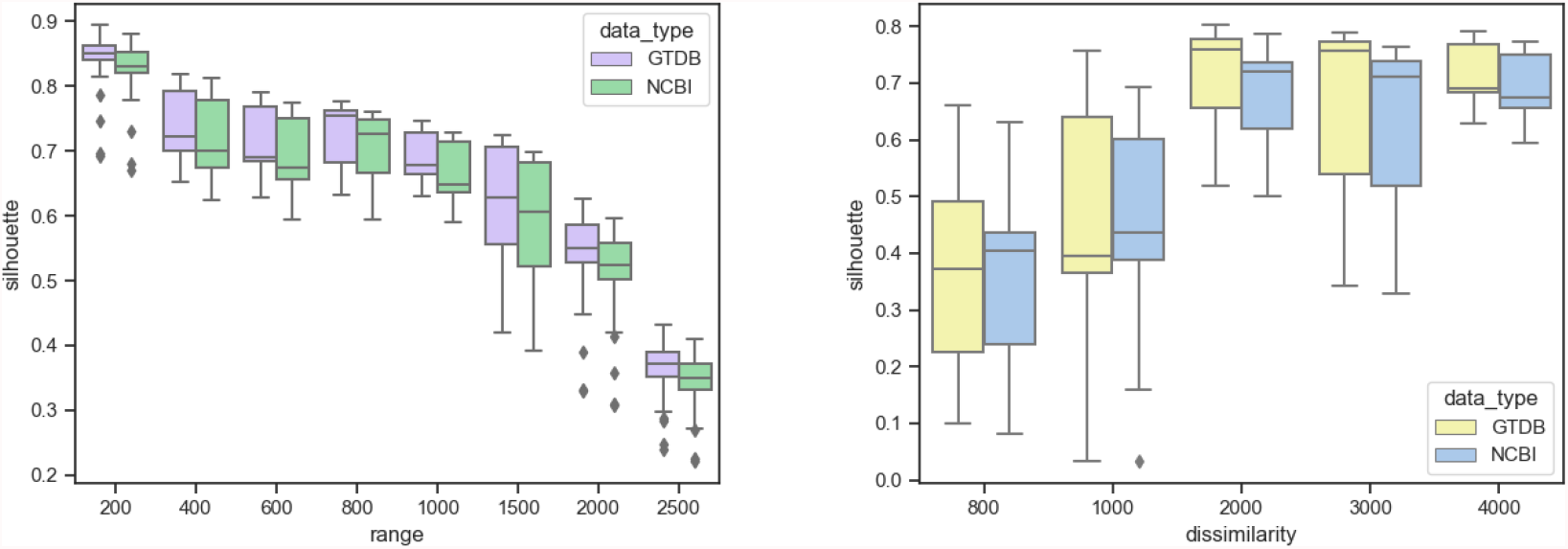
A comparison between the Silhouette scores computed using the GTDB taxonomy and NCBI taxonomy.

Even with differing underlying taxonomic tree topology, we observed highly similar behavior of UniFrac when using the GTDB taxonomy and when using the NCBI taxonomy. In some cases, specifically, when dissimilarity was relatively small, the NCBI taxonomy appeared to yield slightly better performance when WGSUniFrac was applied. In most other cases, GTDB taxonomy yielded slightly better overall results, which agreed with previous experiments where 16S data yielded better overall results. This is due to GTDB taxonomy being more consistent with 16S-derived taxonomy compared to the NCBI taxonomy. Nonetheless, the similarity in performance between the approaches using the GTDB taxonomy and the NCBI taxonomy, together with the previous experiment, suggest that neither the granularity of the branch lengths nor the taxonomic topology is a significant limiting factor to the application of UniFrac, supporting our hypothesis.

### 2.3 On simulated reads

In the previous section, it has been demonstrated that WGSUniFrac is able to cluster samples according to environments in the most ideal situation in which both the 16S OTU tables and WGS profiles were created without the consideration of sequencing errors and profiling biases, which are common in real-world applications. In addition, different profiling methods and taxonomic classification methods may produce different results both between 16S and WGS data and within the same data type [20, 34, 44].

To answer the question if WGSUniFrac would remain robust under a more realistic setting, in this section we investigate the performance of WGSUniFrac on profiles produced from simulated reads. We also increased the complexity of the experimental setup by testing not only with two environments but also with five.

We used Grinder [3] to simulate both 16S amplicon reads and WGS reads with sequencing protocols similar to those of common modern-day sequencing platforms as much as possible while maintaining computation efficiency (see Supplementary Experimental setup details). We used the built-in Dada2 [6] plugin in QIIME [7] to infer taxonomic feature tables from 16S amplicon reads. We used mOTUs [35] to generate taxonomic profiles from the simulated WGS reads. We then calculated and compared UniFrac and WGSUniFrac respectively on the results.

Following a similar approach as section 2.1, the following setups were conducted twice, one using two environments and the other using five: Fixing the range to be 500, we generated experiments having dissimilarities 1,000 to 6,000 in steps of 1,000; fixing dissimilarity to be 4,000, generate experiments with range 200, 1,000, 2,000, 3,000. Each of these combinations was repeated five times with organisms chosen at random. The results are summarized in Table 1 and Figure 6.

**Table 1:** Mean Silhouette Indices for 16S and WGS clusterings by pairwise UniFrac. Higher Silhouette index indicates better clustering of environments.

|  | 2 Environments | 5 Environments |
| --- | --- | --- |
| 16S | 0.226 | 0.051 |
| WGS | 0.562 | 0.206 |

**Figure 6:**
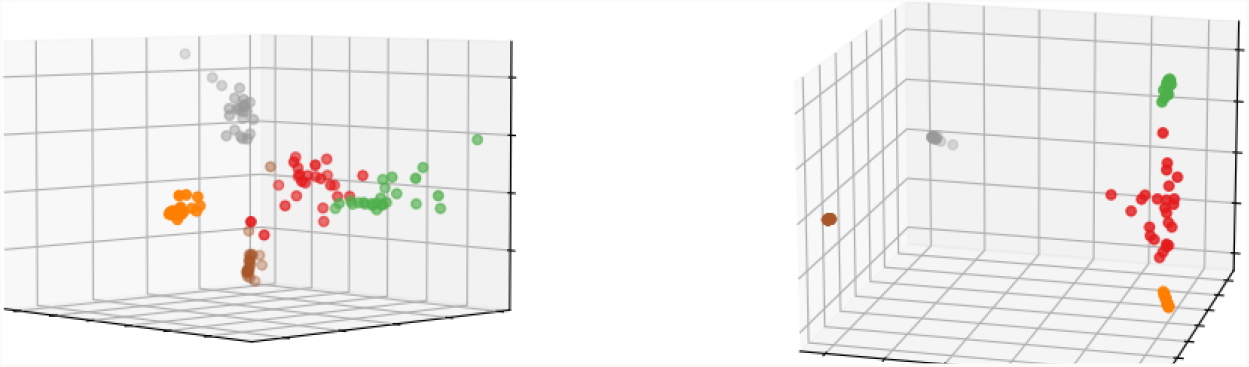
An instance of the comparison between PCoA plots produced using 16S and WGS data with range 300 and dissimilarity 4,000, colors depicting environments. Left: 16S UniFrac. Right: WGSUniFrac.

It was somewhat surprising that the mean Silhouette scores significantly favored the WGS approach in contrast to the 16S approach, which was expected to have better performance. This could be due to the intrinsic differences in simulation protocols and tools used. It has also been pointed out that abundance profiling has much better accuracy when WGS data is used compared to when 16S data is used [20]. This might potentially explain the poor performance of 16S data when inferring of abundances from reads was involved, which also shows the limitation of 16S data and motivates our endeavor to explore a good metric that can be applied to WGS data. Still, an average score of 0.562 allowed us to believe that UniFrac can be applied to WGS data even in the presence of sequencing errors and noises.

### 2.4 On real WGS studies

While running the experiment on simulated reads allowed a glimpse of the feasibility and performance of WGSUniFrac in a more realistic setting, the real-world situation is still much more complex. For instance, the organisms involved in the previous experiments all come from one single phylogenetic tree [10, 31]. In each experiment setting, organisms were selected to simulate distinct environments, with each sample consisting of the exact same number of organisms with relative abundances distributed over a near-ideal exponential distribution. Also, in order to have a fair comparison with 16S UniFrac, combined with limitations of tools in read simulation and profiling processes, compromises such as limiting read lengths were made, further impacting the resemblance between the simulated data and potential real world data.

As such, we proceeded to test WGSUniFrac on real world studies using human whole genome shotgun data. It has been observed and reported in various 16S studies that metagenomic samples collected at different body sites of a human significantly differ [9, 17, 21]. We investigated if this property could be captured using WGS data alone by investigating if samples can be clustered depending on the site of collection.

Using the HumanMetagenomeDB database [18], a curated database for human WGS metagenomic data, we searched for metagenomic projects with specified body sites. To minimize the effect of differences in sampling and sequencing protocols in different studies, we limited our search to studies originating from the Sequence Read Archive (SRA), sequenced using ILLUMINA, and with number of sequences 10 million and above. Among these, we considered only paired-end data and applied the same quality control to all samples prior to profiling to maintain consistency across samples as much as possible (See Supplementary Materials Experimental setup details). The samples were then converted to taxonomic profiles using mOTUs [35]. Among these profiles, we removed those containing less than 100 species. The resulting PCoA plots are shown below. To eliminate the potential bias that the samples might be clustering by studies instead of by body sites, as most studies involved one single body site each, we also produced the PCoA plot colored according to project ID for each category as a comparison.

From Figure 7, we can see that samples were clustered with reasonable sensitivity according to body sites rather than by study, despite the varying protocols across studies, a demonstration of the robustness of WGSUniFrac in real-world applications.

**Figure 7:**
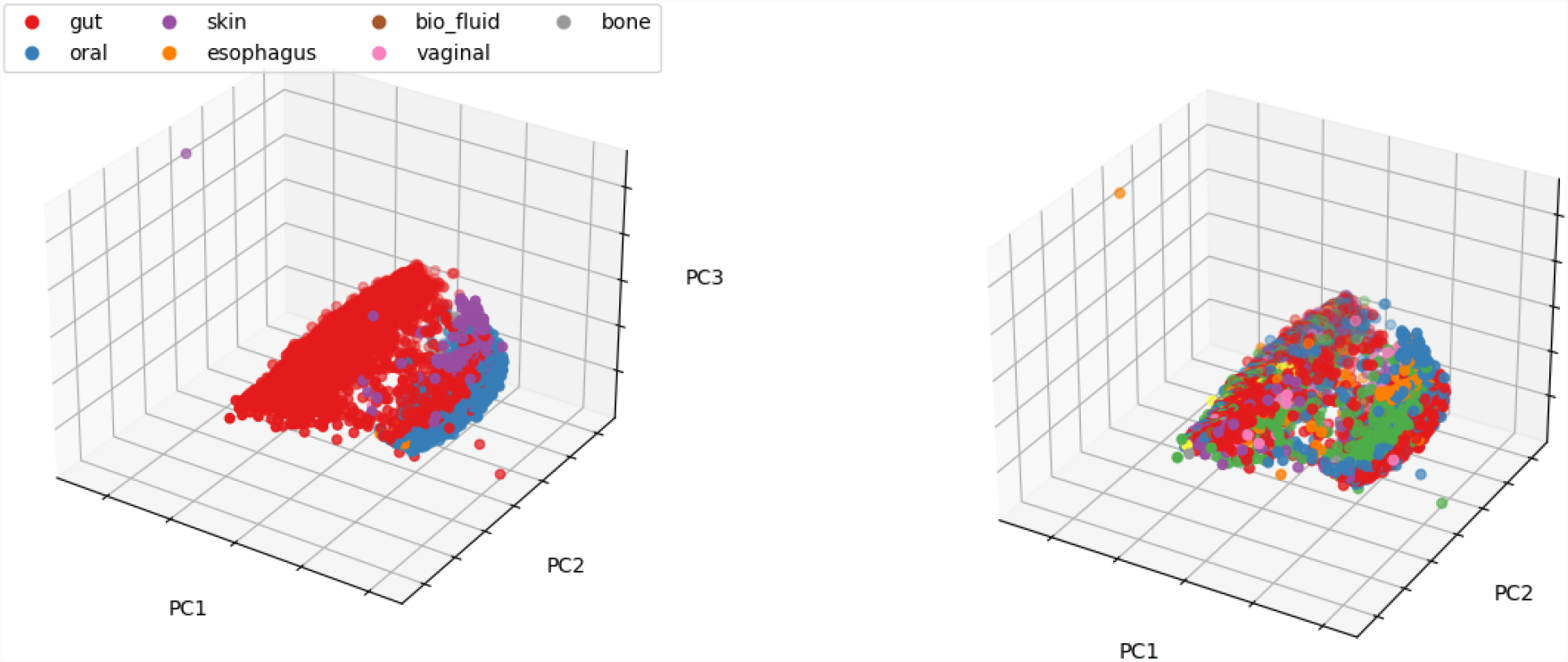
Left: samples colored by body sites. Right: sample colored by studies.

At this point we revisit the open problem of branch lengths function selection in section 1, using these real data. Since the number of data points were massive, for the ease of observing patterns, we stratified the profiles into three categories and analyse them separately: low diversity (containing 100 to 200 species), medium diversity (200 to 300 species), and high diversity (300 species and above). For each of these categories, we produced PCoA plots using branch lengths functions *x*^−1^ and *x*^−2^ respectively. The results are shown in Supplementary Figure S3.

A careful examination of the plots shows that changing the *k* value from -1 to -2 in the branch lengths function *x*^*k*^ only resulted in scaling of the clusters. Specifically, it only clustered more tightly what had already been clustered and revealed no additional information. Hence, there is no strong reason that -2 should be favored over -1. The user could potentially decide on the magnitude of *k* depending on the alpha diversity of the samples, if this information is known.

## 3 Discussions

Up to this point, we have tested the performance of WGSUniFrac in comparison to the traditional UniFrac applied to 16S data under various settings, ranging from the most ideal scenario to real-world data. Under the most ideal scenario, where samples with a phylogenetic classification were directly compared to the corresponding taxonomic classification, WGSUniFrac exhibited comparable ability to distinguish samples from different environments under various parameter settings, providing evidence for the hypothesis that UniFrac can be applied to WGS data simply by assigning branch lengths to a taxonomic tree without significant loss of information on beta-diversity. We then further investigated the effects of different branch length assignments and reached the conclusion that having branch lengths inversely proportional to the height of the taxonomic tree best capitulated the expected clustering trend, while fine-tuning of the magnitude of this proportion did not seem to reveal additional information.

A more detailed investigation of the effect of differences in branch lengths assignments was conducted using the GTDB data, with which we investigated the effect of phylogenetic information both in terms of branch lengths and topology. The results showed that neither the decrease in the resolution of branch lengths nor the change of topology from that of GTDB taxonomy to the conventional NCBI taxonomy significantly decreased the quality of clustering.

The results were slightly puzzling when read simulation was involved in the second part of the experiments, with WGSUniFrac outperforming 16S UniFrac in most cases. We conjecture that this was due to the limitation of simulation and profiling tools and the intrinsic differences in data preparation protocols between 16S and WGS data. The poor performance of 16S UniFrac when sequencing errors were involved demonstrated the potential superiority of WGSUniFrac in real applications. However, further studies are needed to confirm this conjecture. The limitation of efficient read simulation tools that simulate both 16S rRNA and WGS data impeded our further investigation into this matter.

It was perhaps most interesting to evaluate the performance of WGSUniFrac on real data. To this end we tested the ability of WGSUniFrac in recapitulating a known phenomenon previously demonstrated by UniFrac applied on 16S data. Though the lack of corresponding 16S counterparts made a direct comparison to 16S UniFrac unpractical, the PCoA plots did clearly demonstrated the ability of WGSUniFrac in clustering metagenomic samples according to body sites, confirming also in this process that the the differences among samples from different body sites are more prominent than the differences of the same body sites across individuals.

It is also noteworthy that except the last experiment where observations were made purely on WGS data without a quantitative or qualitative “ground truth” to compare to, most of the experiments used 16S data as a reference of comparison. However, this was simply because the UniFrac metric was originally designed to be used data with phylogenetic information, which was typically available when 16S data is employed, not necessarily that the 16S phylogeny is indeed the gold standard. In fact, limitations of 16S data in taxonomic classification have been reported in studies [20, 42]. Which undermines the use of 16S as the standard reference. In addition, such as in the case of GTDB, there have been methods capable of producing phylogenetically consistent taxonomy, and has been shown in the experiments above to yield better results than taxonomy without the additional phylogenetic information. This shows that WGSUniFrac will prove itself to be increasingly useful as better methods to uncover the “real” taxonomic classification in WGS data emerges.

## 4 Methods

The UniFrac was first defined in 2005 by Lozupone et al as the fraction of branch lengths unique to only one of the two communities being compared on a phylogenetic tree [26]. This original version of UniFrac (also known as the unweighted UniFrac) is a qualitative measure that decides if two communities differ significantly based on if the computed UniFrac is greater than what would be expected by chance [26]. The weighted UniFrac was introduced soon after to offer insights to the degree of differences by taking into consideration the relative abundances of the organisms [28], and the original computation is given by:

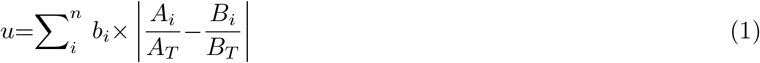

Where *n* is the total number of branches on the tree, *b*_*i*_ is the length of branch *i, A*_*i*_ and *B*_*i*_ represent the number of sequences descended from branch *i* in communities *A* and *B* respectively, and *A*_*T*_ and *B*_*T*_ are the respective total number of sequences for the purpose of normalizing the abundances in the case of uneven sample sizes for communities *A* and *B* [28]. The original UniFrac was only intended for an application on phylogenetic trees reflecting the evolutionary relationship amongst the organisms and on which all the abundances are found on the leaf nodes.

In a previous study, it has been demonstrated that the weighted UniFrac distance is equivalent to the Kantorovich-Rubinstein metric, also known as the earth mover’s distance [12]. Under this definition, instead of building a phylogenetic tree from scratch using the samples, a pre-existing reference tree can be used [12]. By mapping the reads to the appropriate nodes on the reference tree through comparative methods, the information of relative abundances gets incorporated into the tree. The equivalence with the earth mover’s distance then allows us to view the UniFrac distance in a new light: viewing the relative abundances as piles of sand, the UniFrac can be defined as the minimum amount of work required to move the sand from the configuration of one sample to match that of the other, with the amount of work being defined as mass multiplied by the total distance traveled along the tree branches [12]. This gives us an alternative formulation of UniFrac which will be described below.

Let *T* be a rooted tree with *n* nodes ordered from leaves to the root *ρ* representing organisms and branch lengths proportional to evolutionary distances. For a node *i* in *T*, define depth(*i*) as the number of branches on the shortest path from *i* to the root node. We impose a partial ordering on the set of all nodes in *T* in terms of depth: a node *i* is below a node *j* if depth(*i*) *>* depth(*j*). Represent a branch length by *l*(*i*), indicating the weight on the branch connecting node *i* to its ancestor *a*(*i*). Let *P* and *Q* be vectors of probability distribution on the tree with non-negative entries summing up to 1, representing the relative abundance of each organism on the tree in the two input samples respectively, ordered from leaves to the root. Given a node *i* in *T*, let *T*_*i*_ be a subtree of *T* not containing *ρ* obtained by deleting (*i, a*(*i*)). Define *w*_*i*_ to be an indicator function that represents a subtree rooted at node *i* such that the *j*-th entry of *w*_*i*_ equals 1 if (*j, a*(*j*)) is a node in the subtree rooted at *i*, and 0 otherwise. I.e.

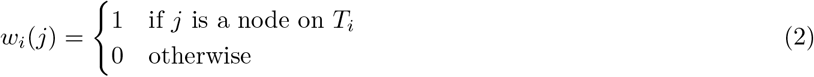

Now let *W* be an *n* × n matrix with column *i* given by *w*_*i*_ and each row *j* scaled by *l*(*j, a*(*j*)). The UniFrac distance (1) can then be represented equivalently as [29, Lemma 2.2.1], [33, Suppl. pg 10]

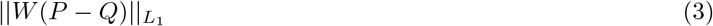

This formulation not only allows the exact UniFrac distance to be computed in linear time [30] but also allows UniFrac to be computed on any tree, not necessarily a phylogenetic one. This allows us to draw one step closer to the application of UniFrac on WGS data, with which a phylogenetic tree is in general impossible to be built, but a taxonomic tree instead. The only obstacle of a direct application lies in the absence of branch lengths *l*(*i*) on taxonomic trees. As a solution we incorporate the assignment of branch lengths according to a given branch lengths function into the algorithm of WGSUniFrac (Algorithm 1) prior to the computation of UniFrac with the EMDUniFrac implementation.

In general, taxonomic trees do not have a natural notion of “branch lengths” as in a phylogenetic tree. As such, we can impose a functional form for the branch *l*(*i*) = *f* (*i, a*(*i*)) where *f* (*i, a*(*i*)) is some function that maps nodes to branch lengths based on some biologically reasonable form. For example, in the Results section above, we chose *f* (*i, a*(*i*)) := (*i*)^*k*^ for *k* ∈ ℤ. Defining *f* in this way means branch lengths are assigned uniformly at each depth, with lengths increasing (or decreasing, depending on the sign of *k*) the further the branches are from the root. The exploration of other values of *k* and their impact on the performance of WGSUniFrac can be found under the Results section. One can also imagine a data-derived definition of the branch lengths if given access to, say, the rate of accumulation of mutations for an organism belonging to the clade defined by the node *i*. In this exposition, the exact form of *f* does not impact the algorithm we describe.

We now give a complete description of the WGSUniFrac algorithm below. Given a rooted tree *T* with nodes ordered from leaves to the root, represented by an edge set *E* = {(*i, a*(*i*))} for *i* ∈ *T*, with *a*(*i*) being the ancestor of node *i*; probability distribution vectors *P* and *Q* representing relative abundances in two samples respectively. For *i* ∈ *T*, let *l*(*i*) = *f* (*i, a*(*i*)) for some function *f* which the user specifies.

### Algorithm 1 WGSUniFrac

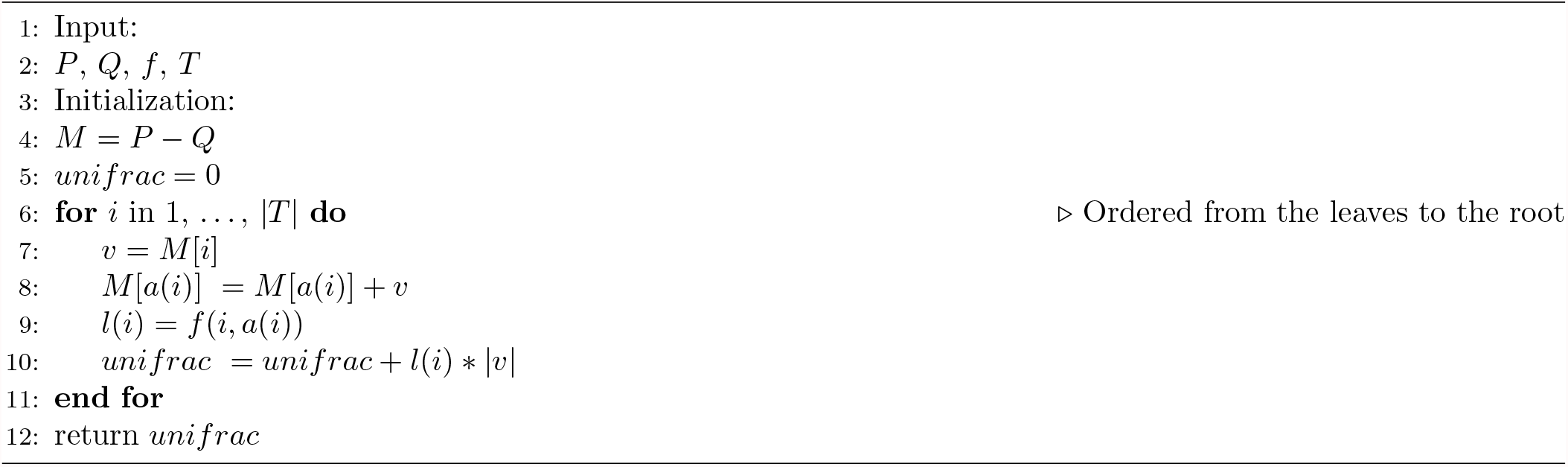

We also give a simple proof that this algorithm does indeed calculate the UniFrac as formulated in equation 3.

Consider the matrix *W* in 3. Let *L* be a vector with the *i*th entry being *l*(*i*) and 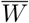 be the skeleton matrix of *W* such that 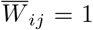 if *W*_*ij*_ ≠ 0 and 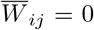 otherwise. Also, for simplicity of comparison, let *M* = *P* − *Q* as in the algorithm. With these notations, (1) can be rewritten as 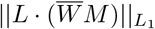 (· denotes the dot product).

By the construction of 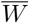, for a given row *i*, 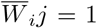 if and only if *j* = *i* or node *j* is an ancestor of *i* on the tree. It is then easy to observe that line 7-9 of Algorithm 1 computes 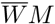. The scaling of 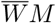 by taking the dot product with *L*, followed by computing the *L*_1_ distance, is done in line 10.

## Supplemental Materials

### S1 Experimental setup details

All computations of UniFrac of 16S data were done using the “beta-phylogenetics” function in `Qiime2` [4]. All profiling of WGS reads into profiles were performed using `mOTUs2` [11] with the parameter “precision”.

#### S1.1 On taxonomic data converted from phylogenetic data

We used the `99_otus` dataset from the `gg_13_5_otus` data package downloaded from Greengenes database [6]. We converted the phylogenetic tree into its corresponding taxonomic tree by using the mapping file provided in the `ete3` python package [7] that maps OTUs to taxonomic IDs and the taxonomic lineage provided in NCBI Taxonomy database. We considered only OTUs in the `99_otus` phylogenetic tree that map to taxonomic IDs with a complete lineage of the ranks superkingdom, kingdom, phylum, class, order, family, genus, and species as can be retrieved from the NCBI Taxonomy database. There are 35,461 such OTUs in total. With respect to each of these OTUs, we computed its phylogenetic distance on the tree (using the “get distance” method in the `ete3` module) from all the other OTUs and obtained a list of OTUs ranked by proximity.

For each experiment, we artificially created two “environments,” each with 25 samples and each containing 200 distinct organisms. The two “environments” were created by first selecting two OTUs sufficiently far apart and each sample was populated by randomly selecting 200 organisms with exponentially distributed abundances among OTUs within a certain range of distance from the two selected OTUs. For simplicity of computation, we used the relative position in the proximity list with respect to an OTU as a proxy to the actual distance between the OTUs. i.e. instead of selecting 200 OTUs among those within a distance of *x* from a given OTU, we choose 200 among *y* OTUs closest to the given OTU. The parameter of the value *y* will be defined as “range” in this paper, whereas the initial “distance”-(number of OTUs apart) between the first two “environment”-defining OTUs, dissimilarity. Each experiment was repeated 100 times.

We evaluated the performance of our method in identifying and clustering the two “environments” using taxonomic (WGS) data in comparison to the performance of traditional weighted UniFrac on its 16s counterpart under various settings of range and dissimilarity.

#### S1.2 Comparison with phylogeny-aware taxonomy

The GTDB data were obtained from https://data.gtdb.ecogenomic.org with release 202. We obtained the `bac120` taxonomic tree together with the corresponding taxonomy. The taxonomic ID for each of the organism in the `bac120` taxonomy file was retrieved using `TaxonKit` [12]. Species without a matching taxonomic ID in any part of the lineage were removed. There were approximately 4,900 species remaining after this process. The general approach for this part of the experiment is highly similar to that of the section above, with the original GTDB tree playing the role of the phylogenetic tree. There were 100 repeats for each combination of range and dissimilarity shown.

For the first part of the experiment, we selected species from the `bac120` taxonomic tree according to the protocols above and treated these samples as 16S samples, computing pairwise UniFrac distance matrix using `Qiime2` [4]. For each sample, its corresponding taxonomic profile was generated, following the GTDB taxonomy as provided in the taxonomy file obtained from the database. The UniFrac distance matrix for each sample was computed using our method with the branch length function *l*(*x*) = *x*^−1^, where *x* is the depth of the tree a branch belongs to, counted from the root.

For the second part of the experiment, the profiles using GTDB taxonomy were used as a reference. For each of these profiles, the species were singled out and for each species, the taxonomic path was reconstructed by retrieving the lineage from NCBI using the `ete3` python package [7], thus creating a second set of profiles differing from the first set only in taxonomic path. The UniFrac matrices of these GTDB profiles and NCBI profiles were compared, using the same branch length function of *l*(*x*) = *x*^−1^, such that the differences in the results were solely accountable by the difference in taxonomy and nothing else.

#### S1.3 On simulated reads

To evaluate the applicability of UniFrac on more realistic data, we tested our method on simulated reads. Both simulations of 16S amplicon libraries and of WGS libraries were done using `Grinder` [2]. For the 16S part, we used the reference genomes `99_otus.fasta` provided in the same `gg_13_5_otus` package from Greengenes as the first part of the experiment. With the aid of the mapping file provided that maps OTUs to NCBI accessions, we used the `esearch` and `efetch` functions in `Entrez Direct` [9] to extract the whole genome of each organism present in the 16S reference genomes, if it existed. To simulate amplicon sequencing reads, we use the forward primer sequence AAACTYAAAKGAATTGRCG as suggested by `Grinder`. Both the amplicon sequencing and WGS sequencing were single-end, with read length 150bp, 4th degree polynomial error model parameters suggested by `Grinder` and the default 80:20 substitution:indel error ratio, 5× coverage for 16S reads and a total read number of 1,000,000 for WGS reads.

The resulting 16S libraries were denoised using `Qiime2` plugin `dada2` [3], with phylogenetic tree built using `Qiime2` plugin fragment-insertion `SEPP` method [8], and finally converted to pairwise UniFrac distance matrix. The WGS libraries were profiled using `mOTUs` [11] into CAMI format [1] profiles, from which the pairwise UniFrac matrix was computed for each experiment.

Using the same protocol in “environment” creation as the first part of the study described above, with the restriction to only organisms with an WGS reference sequence available on NCBI (around 6000 in total), we simulated either two or five environments for each experiment with varying combinations of range and dissimilarity. Each experiment was repeated five times.

#### S1.4 On real-world studies

We used the HumanMetagenomeDB [10] to filter and select human whole genome shotgun SRA data from nine body parts with number of sequences within 10 to 437 (maximum number in the HumanMetagenomeDB) million and sequenced using `Illumina`, which came out to be 12,261 in total. Among them, we selected only studies that were paired-end. For these paired-end reads, we performed a quality control using `fastp` [5], after which each sample was profiled using `mOTUs` [11]. Among the profiles we removed those having too few species (less than 100).

## S2 Supplementary Figures

**Figure S1:**
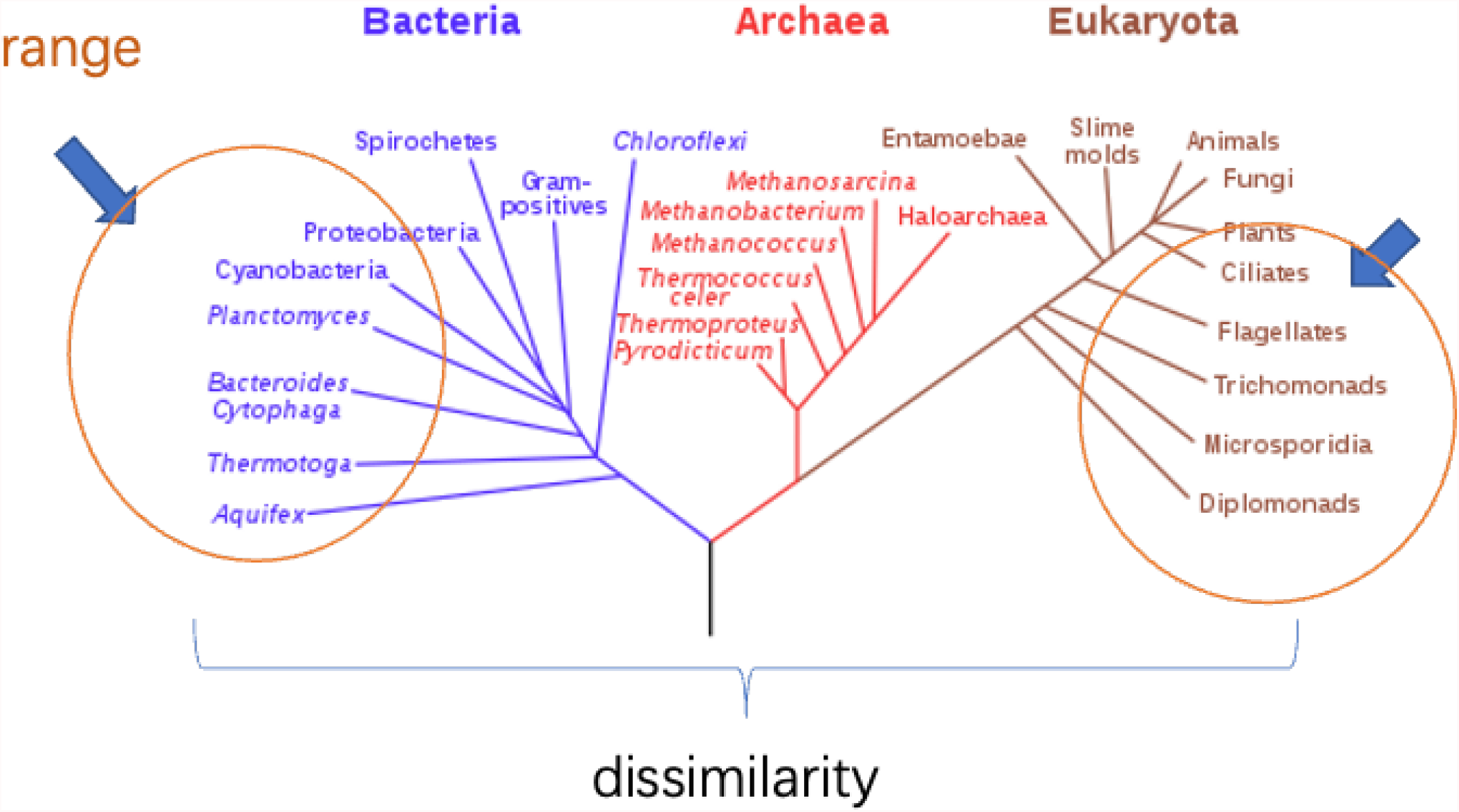
An illustration demonstration the definitions of range and dissimilarity.

**Figure S2:**
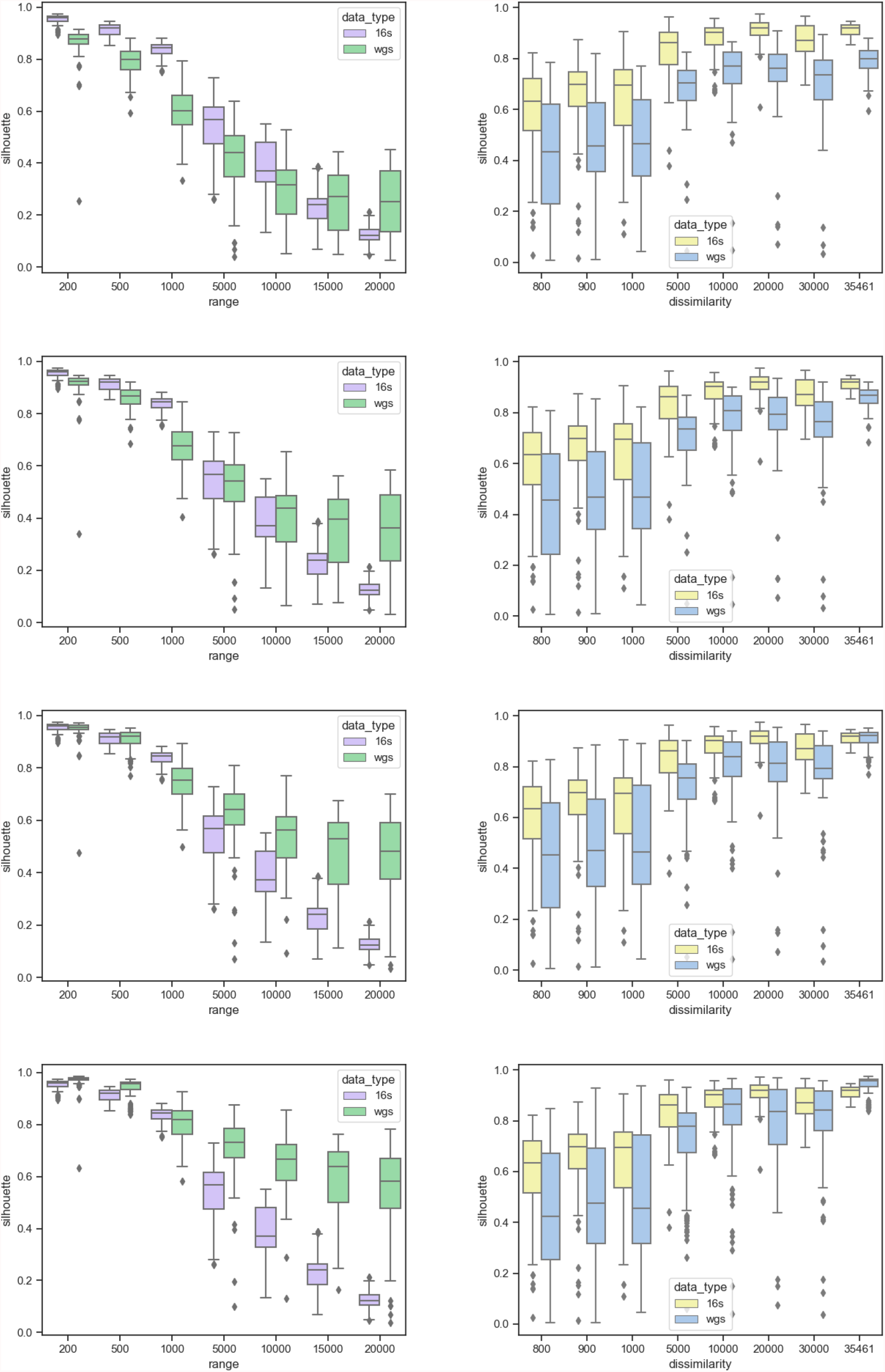
Top panel: Silhouette score against range. Bottom panel: Silhouette score against dissmilarity. From top to bottom: *k* = −0.5, −1, −1.5, −2 respectively.

**Figure S3:**
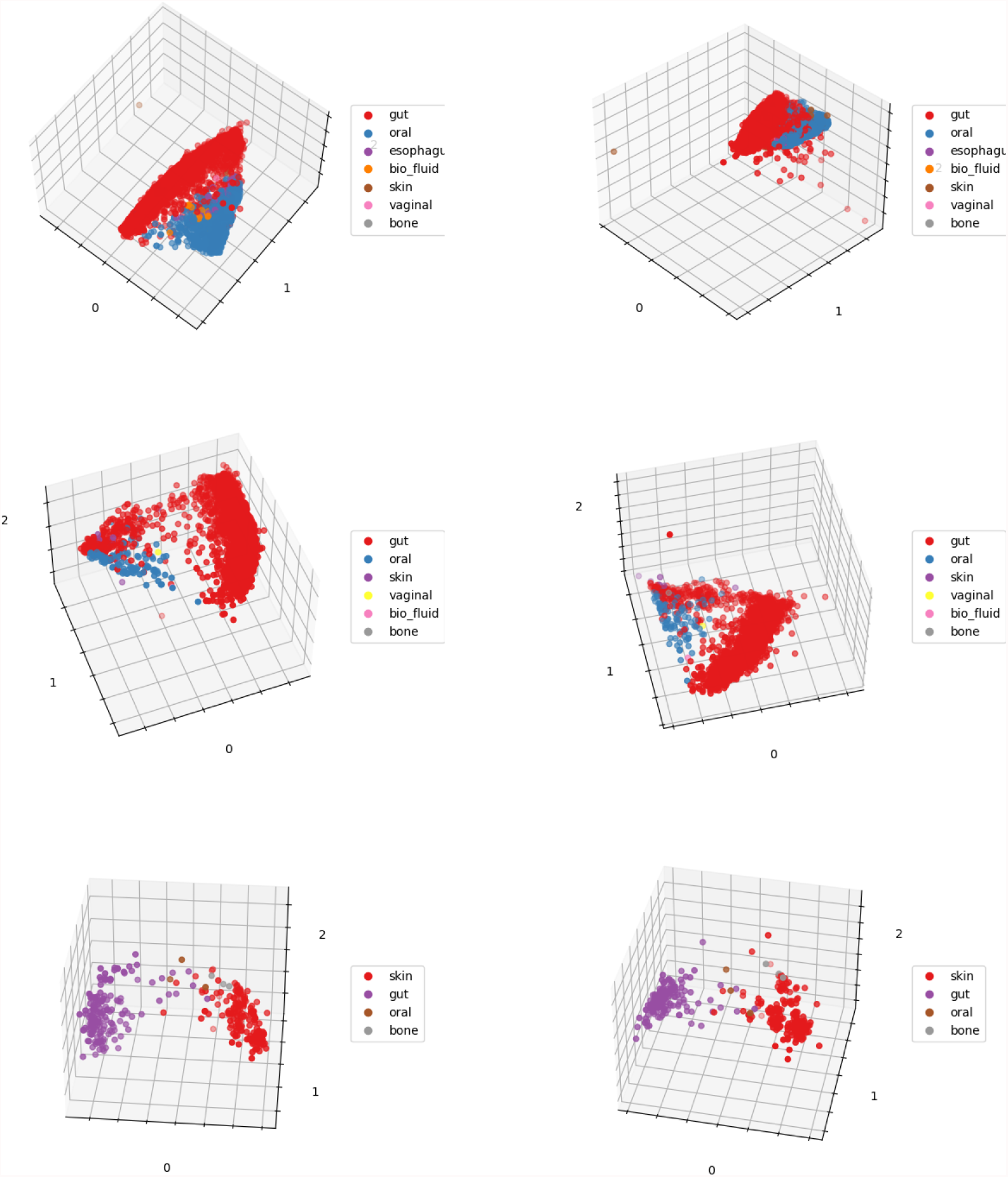
From top to bottom: low diversity, medium diversity, high diversity. Left: branch lengths function *x*^−1^, right: branch lengths function *x*^−2^

**Figure S4:**
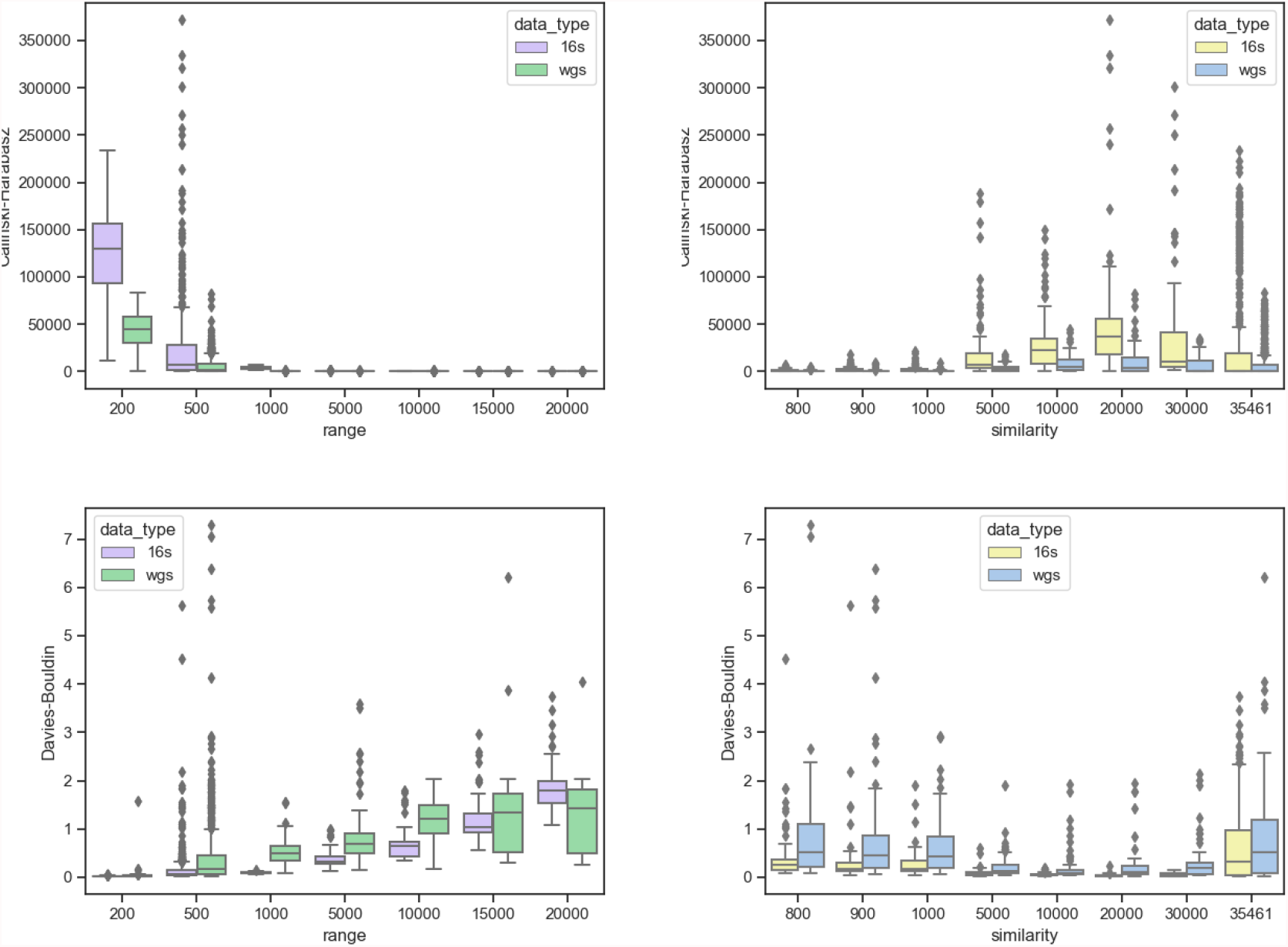
Clustering quality measured using different metrics. Top panel: Calinski-Harabasz index. Bottom panel: Davies-Bouldin index.

## Notes

### Competing Interest Statement

The authors have declared no competing interest.

